# Mutational and biophysical robustness in a pre-stabilized monobody

**DOI:** 10.1101/2020.12.14.422768

**Authors:** Peter G. Chandler, Li Lynn Tan, Benjamin T. Porebski, James S. Green, Blake T. Riley, Sebastian S. Broendum, David E. Hoke, Robert J. Falconer, Trent P. Munro, Malcolm Buckle, Colin J. Jackson, Ashley M. Buckle

## Abstract

The fibronectin type III (FN3) monobody domain is a promising non-antibody scaffold which features a less complex architecture than an antibody while maintaining analogous binding loops. We previously developed FN3Con, a hyper-stable monobody derivative with diagnostic and therapeutic potential. Pre-stabilization of the scaffold mitigates the stability-function trade-off commonly associated with evolving a protein domain towards biological activity. Here, we aimed to examine if the FN3Con monobody could take on antibody-like binding to therapeutic targets, while retaining its extreme stability. We targeted the first of the Adnectin derivative of monobodies to reach clinical trials, which was engineered by directed evolution for binding to the therapeutic target VEGFR2; however, this function was gained at the expense of large losses in thermostability and increased oligomerisation. In order to mitigate these losses, we grafted the binding loops from Adnectin-anti-VEGFR2 (CT-322) onto the pre-stabilized FN3Con scaffold to produce a domain that successfully bound with high affinity to the therapeutic target VEGFR2. This FN3Con-anti-VEGFR2 construct also maintains high thermostability, including remarkable long-term stability, retaining binding activity after 2 years of storage at 36 °C. Further investigations into buffer excipients doubled the presence of monomeric monobody in accelerated stability trials. These data suggest that loop grafting onto a pre-stabilized scaffold is a viable strategy for the development of monobody domains with desirable biophysical characteristics, and is therefore well-suited to applications such as the evolution of multiple paratopes or shelf-stable diagnostics and therapeutics.

## Introduction

Developing a small, simple protein domain that includes a similar sized binding region to antibody complementarity determining regions (CDRs) is a successful strategy for overcoming the complexity of antibody structure. Non-antibody scaffolds are single domains, typically smaller than 20 kDa in molecular weight, and mostly free of glycosylation or disulfide bonds that require eukaryotic expression (1). Critically, they exhibit comparable binding affinities to antibodies (2). Monobodies based on the Fibronectin Type 3 domain (FN3) are a popular scaffold for developing non-antibody therapeutics (2– 5). The FN3 domain has an Ig-like fold and thus retains three of the CDR-like loops of an antibody variable fragment, but is structurally simple enough to be engineered for advanced non-antibody functions as the monobody scaffold (4–7). There are a number of unique monobody derivates in active development including clinical Adnectins by LIB therapeutics (8) or ViiV Healthcare (9), the stability-enhanced Centyrins under ARO therapeutics (10, 11) or in a CAR-T format by Poseida Therapeutics (12), and also the related TN3 monobodies under Viela Bio (13).

An important consideration in the ability to evolve a non-antibody scaffold for binding is the combination of a high initial stability and a mutationally robust framework. The small size and lack of redundant framework regions in non-antibody scaffolds results in protein domains which will accumulate only a few mutations to their variable regions before stability becomes compromised (2, 14). Most monobody derivatives typically lose ∼40 °C of thermostability upon evolution for binding (2) which often results in insoluble expression in bacteria that must be resolved through later rounds of evolution (15). Critical to the design of next generation therapeutics, poor biophysical properties such as thermostability and poor or insoluble expression hinder scaffold ‘developability’ and correlate to higher risk of failure in during clinical development (16, 17).

There are multiple approaches of improving the stability of protein folds (18). Protein stability exists between two critical thresholds, where a protein is able to fold into a stable, native 3-dimensional shape but is also still dynamic enough to perform its functions. As proteins evolve over time, the resulting mutations are more likely to be detrimental to stability of the native fold than neutral or positive, and reduce protein stability below the folding threshold (19). This is often considered as a natural trade-off between stability and activity, and can be a severe limitation to directed evolution experiments (20, 21). Therefore, as a protein scaffold takes on mutations to a variable region to improve binding, it is at risk of deleterious losses in thermostability. Accordingly, proteins with improved initial thermostability are able to sample a larger proportion of this destabilizing sequence space as neutral mutations – they are more *mutationally robust* – which in turn increases the chance of reaching novel functions (22–24). In this way, pre-stabilization can enhance both evolvability - the ability of a protein to evolve new functions (21), and developability - the biophysical likelihood of successful development from a lead protein into a therapeutic drug (17).

From a drug development perspective, the native thermostability of a protein correlates with expression titre and improves critical quality attributes such as shelf-life (16), while also expanding the range of storage formulations which can be used in a drug product (25). Biological formulations need to remain stable for at least 2 years at 5 °C, and storage buffers are usually applied to reach this target. This has resulted in a standard set of buffer formulations across industry that are well-validated and focused on reducing aggregation or other loss of active protein (26). However, this sole focus on protein stability limits the exploration of non-standard formulations which benefit other critical quality attributes (25), such as controlling viscosity or osmolarity to lower the pain associated with injection (27–29).

Multiple, well-established techniques use phylogenetic information to derive thermostable homologs for a given protein family (18, 30–34). Here we have specifically applied consensus design, which identifies conserved residues within a protein family that are likely to be stabilizing (31, 34). Previously, we used consensus design to produce the pre-stabilized FN3Con monobody, which has substantially increased thermostability compared to similar fibronectin domains (35). The mutational robustness of FN3Con was then demonstrated by grafting binding loops from an FN3 scaffold that had been selected by laboratory evolution to bind lysozyme with high affinity (36). Whereas binding function was completely transferred, the trade-off in thermostability was negligible compared to that which occurred upon directed evolution of the original scaffold.

In the current study, we aimed to establish whether FN3Con can harbour valuable loop sequences that confer clinical inhibition of a target but were detrimental to stability in established scaffolds. For this study we chose an Adnectin domain that was previously subjected to directed evolution for high affinity binding to the therapeutic target VEGFR2, with function gained at the expense of large losses in thermostability and increased oligomerisation propensity (37, 38). We grafted binding loops from Adnectin-anti-VEGFR2 onto the FN3Con scaffold to produce a recombined domain that retained binding affinity. The FN3Con-anti-VEGFR2 graft was expressed in *E. coli* with little aggregation and maintained characteristically high thermostability, including 24-month stability at 37 °C. An early exploration of buffer excipients produced further stability improvements. We discuss the implications of generating clinical leads by salvaging loop sequences from scaffolds with challenging biophysical features and the importance of designing highly ‘evolvable’ constructs on downstream factors of scaffold ‘developability’.

## Results

### Transfer of affinity to a target by sequence grafting

We chose the Adnectin-anti-VEGFR2 monobody ‘CT-322’ as a candidate for loop grafting to the hyperstable FN3Con in order to test our hypothesis that a stabilized scaffold can rescue stability losses accrued after evolutionary selection for high affinity binding. This Adnectin was generated from mRNA display against a construct of 7 extracellular domains of VEGFR2 fused with a human antibody Fc region, generating high affinity binding (K_D_ = 0.31 nM) but with a 34 °C loss in thermostability (*T*_m_) and also at cost to oligomerisation resistance (37, 38).

Given the absence of structural information for the binding mechanism of Adnectin-anti-VEGFR2 to its large multidomain target, we used previously established loop sequence boundaries (36) to guide the transfer of evolved binding loops to FN3Con, designing FN3Con-anti-VEGFR2 [**Figure 1**A]. Additionally, the entire Adnectin C-terminal tail was reported to be critical to high affinity binding and was also transferred (37).

**Figure 1:**
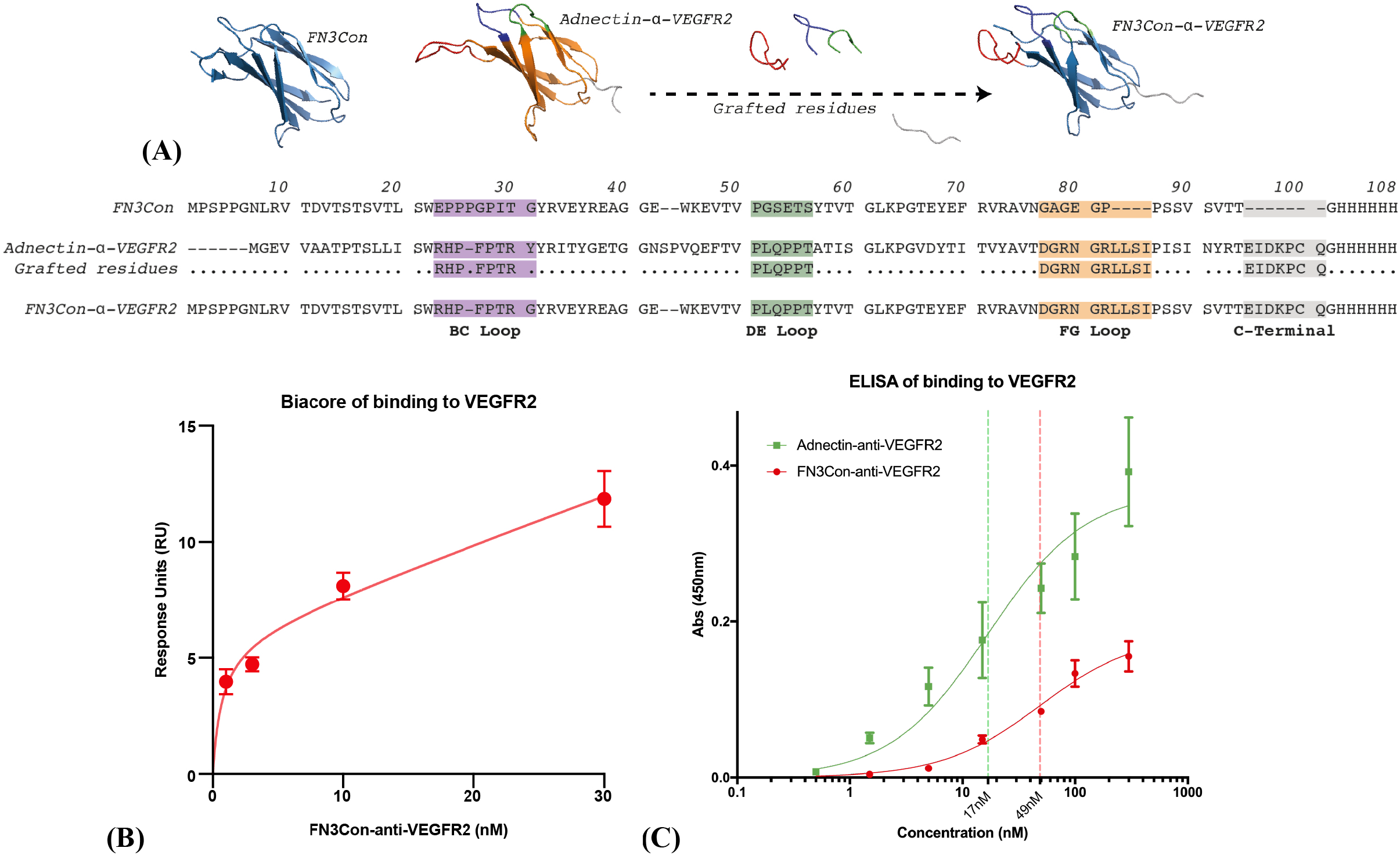
**(A)** Sequence-based grafting between monobody domains with loop sequences from (38). Affinity of FN3Con-anti-VEGFR2 to VEGFR2 was measured using **(B)** Response Units during steady-state of association in Surface Plasmon Resonance (SPR), full data in [**Figure S1**], and **(C)** a general ELISA approach.

FN3Con-anti-VEGFR2 displayed high affinity binding to VEGFR2 (K_D_ = 0.72 nM) [**Table 1**], very similar to that of Adnectin-anti-VEGFR2 (K_D_ = 0.31 nM) (38). We carried out affinity measurement through an orthogonal approach, where two independent methods provided a K_D_ range of 0.72 – 48.79 nM [**Figure 1**B,C **Table 1**], with the K_D_ of 0.72 nM derived from Biacore data presenting the most robust fits to derive underlying equilibrium constants while controlling for confounding non-specific binding and mass transport effects.

**Table 1:** Methodology and results for VEGFR2 binding experiments in Figure 1.

| Protein | Surface Plasmon Resonance (SPR) $K_D$ | ELISA $K_D$ |
| --- | --- | --- |
| Adnectin-anti-VEGFR2 | 0.31 nM * | 16.87 nM |
| FN3Con-anti-VEGFR2 | 0.72 nM | 48.79 nM |
| **Analysis** |  |  |
| Fold Difference | 2.32 | 2.89 |
| FN3Con-anti-VEGFR2 Model Fit | $R^2 = 0.95$ | $R^2 = 0.98$ |
| Immobilised protein | VEGFR2 | VEGFR2 |
| Measure | Change in refractive index | Concentration of Biotinylated protein |

\* SPR $K_D$ from (Mamluk et al., 2010)

### Sequence-based redesign stabilizes an anti-VEGFR2 fibronectin

Adnectin-anti-VEGFR2 undergoes irreversible thermal denaturation with a *T*_m_ of 50±0.4 °C, as measured by circular dichroism (CD), with visible precipitate upon cooling [**Figure 2**A]. In striking contrast, FN3Con-anti-VEGFR2 unfolds reversibly with a *T*_m_ of 89±0.2 °C [**Figure 2**B]. Usefully, the expression yield of FN3Con-anti-VEGFR2 in *E*. Coli increased >10 fold from the parent FN3Con scaffold (not shown). The *T*_m_ of the Adnectin-anti-VEGFR2 was not previously published (38), but our results closely match a precursor Adnectin-anti-VEGFR2 variant of similar affinity, thermostability, and loop sequences (37). This trajectory of loss in Adnectin thermostability presents the trade-offs that take place as affinity is further matured [**Figure 4**A]. In contrast, while the FN3Con scaffold loses ∼10 °C of thermostability after loop-grafting, scaffold stability remains substantially higher than the parent Adnectin molecules, while also retaining reversible refolding.

**Figure 2:**
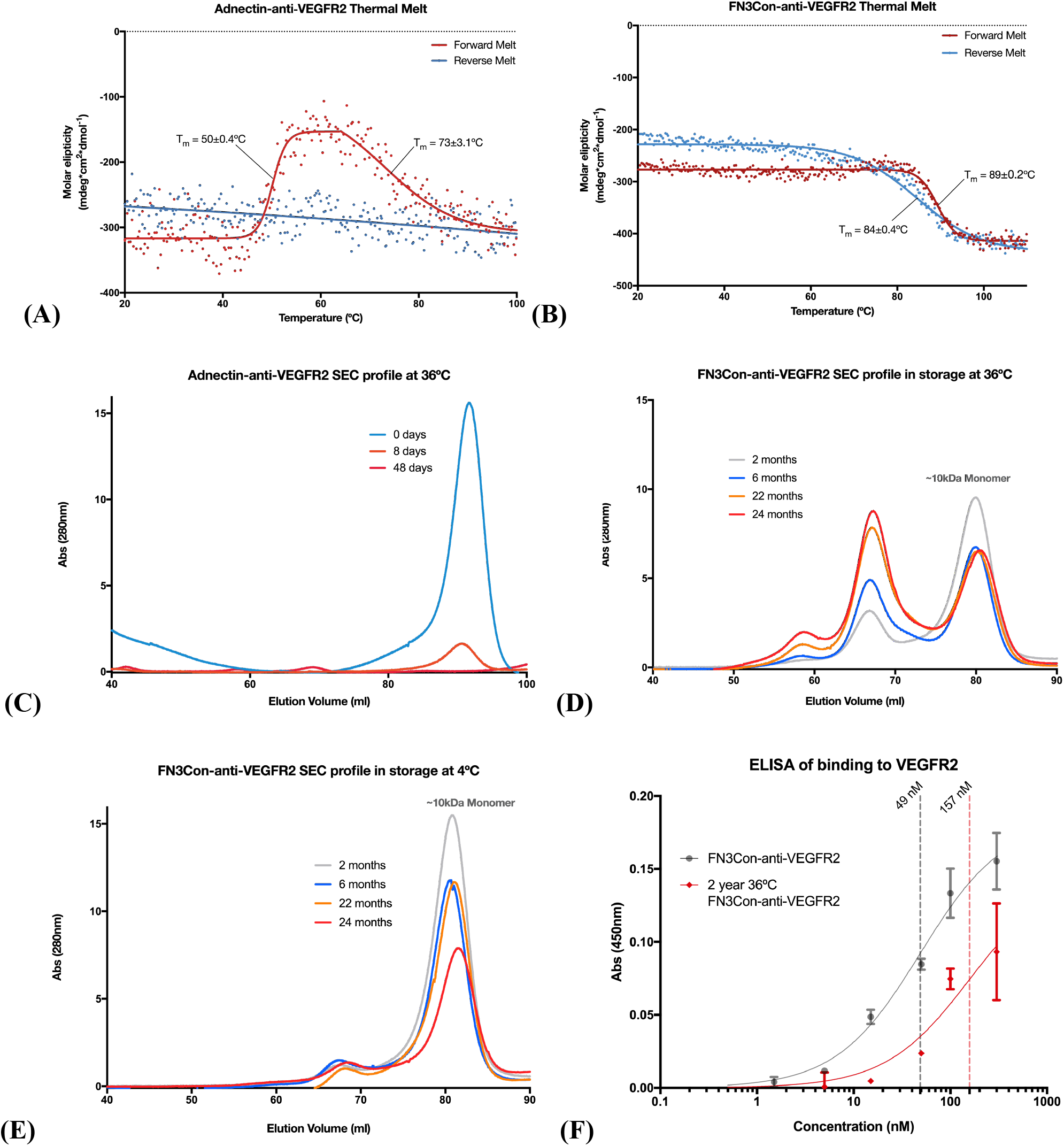
Circular dichroism (CD) thermal melts of **(A)** Adnectin-anti-VEGFR2 (*T*_m_ of 50±0.4 °C) and **(B)** FN3Con-anti-VEGFR2 (*T*_m_ of 89±0.2 °C), a reverse melt (blue) indicates reversible refolding in FN3Con-anti-VEGFR2. **(C)** The Adnectin-anti-VEGFR2 sample aggregates completely over 1 month at 36 °C. **(D)** More than 1/3 of FN3Con-anti-VEGFR2 sample is still present as a monomer after 2 years of 36 °C storage, with the remaining 2/3 present as higher order oligomers. **(E)** The graft also shows limited aggregation during 4 °C storage. **(F)** ELISA of the 36 °C 24-month sample shows a matching 3-fold loss in affinity (K_D_ of 49 – 157 nM).

### FN3Con-anti-VEGFR2 retains stability and function at 36 °C for two years

We next investigated the effect of FN3Con-anti-VEGFR2 hyper-stability on long-term stability and binding activity, as a proxy for extended shelf-life. The Adnectin-anti-VEGFR2 sample completely aggregates within one month’s storage at 36 °C in PBS [**Figure 2**C]. In contrast, the FN3Con-anti-VEGFR2 remained in solution for at least 24 months, with ∼30% of the total sample remaining as a monomer after 2 years storage at 36 °C [**Figure 2**D]. Up to 50% of the FN3Con-anti-VEGFR2 high-order species observed formed between 0 and 6 months at 36 °C, after which oligomer formation stabilized. Accordingly, FN3Con-anti-VEGFR2 presented extended long-term stability at 4 °C, remaining as a monomeric protein in extended trials up to 24 months of storage in PBS buffer [**Figure 2**E]. Strikingly, high affinity binding (K_D_ ∼157 nM) to VEGFR2 was maintained after 24 months at 36 °C [**Figure 2**F], although the observed affinity for the total sample is 3-fold lower than ‘fresh’ FN3Con-anti-VEGFR2 (K_D_ ∼49 nM). This suggests that only the ∼30% monomeric fraction retains binding affinity to the target.

### Matching protein thermal stability with formulation stability

Given the results from accelerated stability testing, our final investigation explored the effect of stabilizing excipients on further improving the shelf-life properties of the FN3Con-anti-VEGFR2 construct. After incubation at 40 °C for 30 days with five different buffer excipients (16); size exclusion chromatography revealed that amino-acid excipients arginine, histidine, glycine and aspartic acid produced a doubling of monomeric FN3Con-anti-VEGFR2 sample [**Figure 3**]. Excipients like Tween80 provide resistance to factors such as hydrophobic unfolding from shaking during storage (25, 39), and was added with no significantly greater detrimental effect on accelerated thermal stability than with PBS buffer alone.

**Figure 3:**
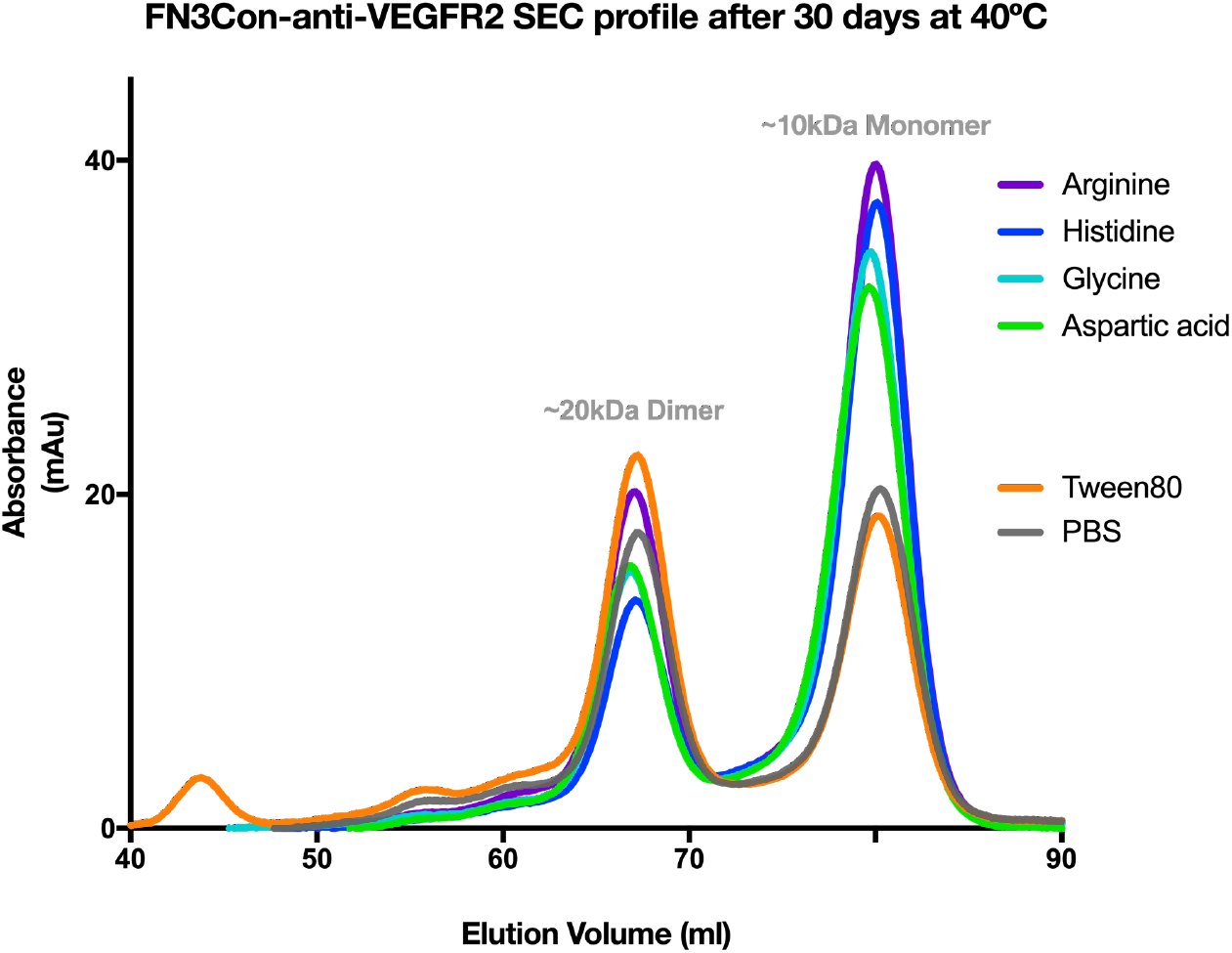
**(A)** SEC elution profiles of FN3Con-anti-VEGFR2 after accelerated stability trials with a range of buffer excipients. All amino acid buffers slowed the loss of monomer. However, dimers appear at a similar rate between all samples, which could be linked to the generation of disulfide-bonded dimers due to degradation of BME. Tween80 was not significantly more destabilizing to the monomer than PBS buffer alone.

## Discussion

Previously, we have shown that FN3Con is robust to the stability-function trade-off (36). The data we present here is consistent with these findings, confirming our hypothesis that a stabilized scaffold can rescue functional regions from the stability losses accrued after evolutionary selection for high affinity binding. Furthermore, our data confirms that grafting onto the FN3Con scaffold improves expression yield (data not shown) and aggregation resistance. Importantly, we show that the stability of FN3Con is suitable for loop grafting to generate high affinity binders against a clinically relevant target.

The Adnectin-anti-VEGFR2 monobody ‘CT-322’ was the first of the Adnectin derivative of monobodies to reach clinical trials (38). Adnectin-anti-VEGFR2 showed potent efficacy with an anti-angiogenic effect in preclinical models of pancreatic cancer (40), colorectal carcinoma and glioblastoma (38, 41), brain tumours (42), and Ewing’s sarcoma (43). Phase 1 clinical studies displayed clinical safety and acceptable pharmacokinetics to support phase 2 studies (44), although only some patients presented stable disease (45).

Unfortunately in phase 2 glioblastoma trials the Adnectin derivative did not produce the required efficacy for continuation of studies (45, 46). This was suspected to be caused by a loss of inhibition effect during translation of this drug from the preclinical setting to human trials (46), which could result from pharmacokinetic issues with the Adnectin scaffold. Related to this could be the emergence of anti-Adnectin antibodies in 31 of 38 patients in the phase 1 study (44, 45). If this failure of clinical translation is driven by issues of developability, such as oligomerisation, immunogenicity and pharmacokinetics, then protein stabilization may salvage a potentially valuable clinical inhibitor.

An underlying explanation for the biophysical limitations of Adnectin-anti-VEGFR2 is that the evolution for high affinity binding came at a critical cost to stability (*T*_m_ decrease of 30 °C) and oligomerisation resistance [**Figure 2**C] (38). The evolution of high-affinity VEGFR2 binding in the Adnectin scaffold resulted in a step-wise trade-off of thermostability for binding over a range of precursor loop sequences [**Figure 4**A] (37, 47). Despite this clinical failure, recent studies have sought to improve the Adnectin’s pharmacokinetic properties by increasing monomer size with a Proline-Alanine-Serine repeat sequence (PAS-ylation) (48). Our loop grafting study examined the ability of FN3Con to mimic the targeted binding of antibodies, supporting the assertion that consensus design of FN3Con resulted in a fibronectin domain that is robust to the mutational load of evolution for function.

**Figure 4:**
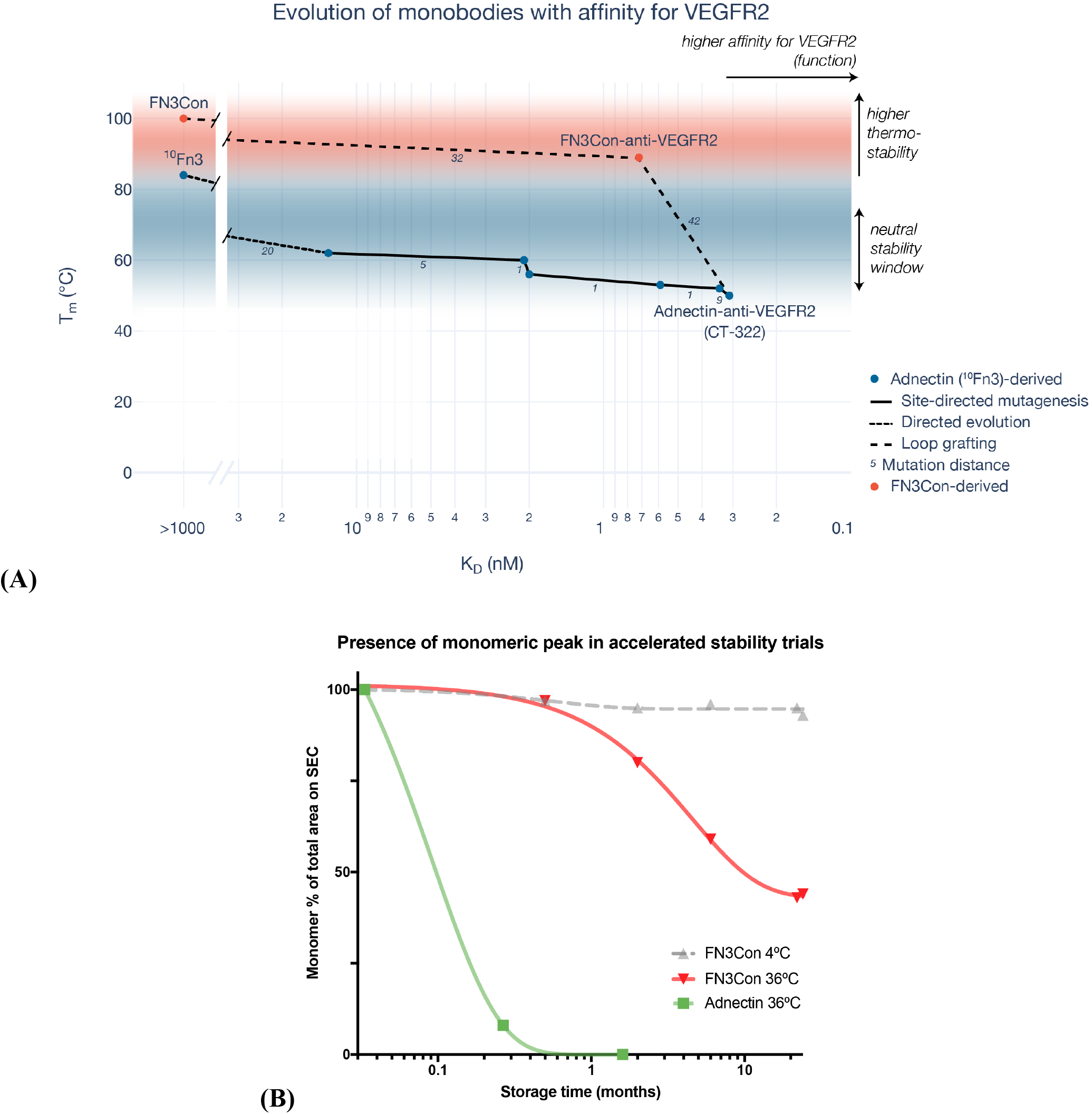
**(A)** Change in protein *T*_m_ as loops of the Adnectin scaffold were evolved for stronger binding to VEGFR2 (37, 38), transfer of binding loops to FN3Con retained similar affinity while heavily improving thermostability. Data in [**Table S1**] **(B)** This improvement in thermal stability produced a related increase in long-term stability under elevated temperatures. The Adnectin aggregated with no stable oligomers, so later monomer percentages were compared against the day 1 values (green). As FN3Con remained at ∼1 mg/ml over the course of each trial, % monomer percentages were calculated from the total peak area within each timepoint (red, gray).

To design FN3Con-anti-VEGFR2 we used available sequence information from the generation of Adnectin-anti-VEGFR2. There is a historic body of work related to rationally grafting CDR loops between antibodies (49). In practice, this grafting technique involves a balance of transferring residues involved in binding and removing stabilizing residues in the accepting scaffold (18, 50), which may reveal certain pairs of donor and recipient scaffolds to be graft-incompatible (36). The Adnectin-anti-VEGFR2 binding loops were compatible with the FN3Con scaffold, given the almost-complete transfer of affinity. This supports the approach that loop-grafting to a more stable scaffold by sequence alone can be an effective strategy for the initial transfer of affinity between fibronectin-like domains, followed by focused rounds of redesign (36, 51).

Furthermore, there is growing evidence that monobody scaffolds can take on antibody CDRs and their resulting affinity to clinical targets (52), such as in the transfer of anti-HER2 CDRs (53). However it is unclear how much of the previously published antibody CDR loop sequences can be transferred to monobody domains, especially if it is a complicated process that involves iterative redesign (51). Additionally, if there are already established antibodies in the clinic then there may be little need for an antibody mimic, unless they can provide meaningful advantages such as short half-life for radiolabelled imaging (5, 54, 55). Nevertheless, it may be more efficient instead to evolve new binding loops *de novo* with display technologies (56, 57). If the FN3Con scaffold is hyper-stable against destructive anti-VEGFR2 binding loops, it may be robust to directed evolution for aspects such as even greater affinity or the addition of a second paratope.

In terms of biophysical properties generally considered under the concept of ‘developability’ (16), FN3Con-anti-VEGFR2 initially presented improved features over the Adnectin in terms of thermal stability and high-yield, soluble bacterial expression (not shown). The improved thermal stability of this construct then led to favourable features of accelerated stability (AS) [**Figure 4**B]. Although it is unclear whether higher-order species in the 2 year AS sample occurred due to self-association or to disulfide-bonding between monomers as BME became inactive (58), the reduction in binding affinity to VEGFR2 matched the loss of monomer in the sample. As a result, in this investigation we see the flow-on effects from the focus on thermostability alone to a broader set of biophysical features, and a link between the concepts of evolvability (59) and developability (16).

Similar to our protein design approach, the purposes of excipient design for stabilization are to either reduce deviation from the native protein conformation or protect that state from collision with other molecules (60, 61). As there is a body of evidence around the interaction of salts negatively destabilizing FN3 proteins (62), we explored excipients which complement the enhanced protein stability of FN3Con. In this study, accelerated stability trials confirmed the protective effect of amino acid excipients (26), as the addition of Arginine, Glycine, Aspartic acid and Histidine doubled the amount of monomeric FN3Con-anti-VEGFR2 in accelerated conditions. The dominant stability improvement of Arginine over Glycine [**Figure 3**] could be due to the complex interaction of the Arginine guanidinium group (63), which suppresses protein unfolding by increasing the energy barrier between folded native-state and aggregation-prone intermediates (64, 65). This can be seen as a further extension of the consensus protein design approach which initially ‘smoothed out’ this folding landscape and slowed the rate of unfolding in the native FN3Con conformation (35, 66). Histidine also provided improved resistance to thermal aggregation and its effect is well-explored in industry formulations (67).

Generating highly stable biological scaffolds provides greater freedom to use non-standard buffer formulations. As biological therapeutics must display a 5 °C shelf-life of up to two years, storage buffers are often designed with an aim of protection from aggregation or chemical modification of the protein (25). With hyper-stability built into a protein scaffold, formulation scientists can instead focus on excipients which optimise function (61), such as for particulate drug delivery depots that overcome the short pharmacokinetic half-life of these small protein scaffolds, for freeze-dried and aerosolised suspensions, or in antimicrobial preservatives for long-term shelf-stable formulations.

These formulation experiments are important for designing differentiated applications from the dominant antibody scaffold, where the FN3Con domain may instead find its place in niche applications (5). The improved thermostability of the FN3Con monobody could allow for greater control over self-association intermediates in order to create novel drug modalities (5). Furthermore, our demonstration of stability and activity over 1-2 years supports the use of consensus-designed Monobodies in diagnostic settings that cannot utilize cold-chain distribution, such as field applications of microfluidic devices (68) Overall, the results of this study show the flow-on effects of pre-stabilizing a protein scaffold, from a greater robustness to evolutionary mutations, improved biophysical properties for clinical development, and potentially enabling a greater range of biomedical applications.

## Conclusions

The initial aims of this study were to examine whether the engineering for hyper-stability in FN3Con allowed for improved ‘evolvability’ of clinical function in this hyperstable monobody derivative compared to other scaffolds that are progressing through the clinic. The complete transfer of affinity to a target was successful through sequence-based grafting alone, and the 40 °C increase in thermostability between monobody scaffolds suggests that the FN3Con scaffold is amenable to taking on affinity to a therapeutic target. Further to these aims, the connection of high thermal stability to biophysical aspects of ‘developability’ was investigated. Long-term stability trials show an active monomeric fraction after 2 years at 36 °C and accelerated stability trials in a range of standard excipients show that there is much room for improvement in extension of that shelf-life. These are promising results for continuing development of the FN3Con scaffold. In future, critical investigations will need to utilise these advanced biophysical features to create novel constructs that provide differentiated patient benefit such as the evolution of multiple paratopes or in shelf-stable applications.

## Author Contributions

PGC, BTP and AMB designed the study and drafted the paper. PGC and LLT performed data acquisition and interpretation. JSG, BTR, SSB, DEH and MB assisted in data analysis, acquisition, and interpretation. DEH, RJH, TPM, CJJ and AMB contributed to writing and conceptual advances.

## Competing financial interests

The authors declare no competing financial interests.

## Acknowledgements

We would like to thank Neelam Shah and Jackie Wilce for assistance with Biacore experiments.

## Materials and Methods

### Protein expression and purification

Genes encoding Adnectin-anti-VEGFR2 and FN3Con-anti-VEGFR2 were chemically synthesized and provided in a pD444-CH (C-terminal 6x His tag, ampicillin resistance) vector by DNA2.0. The resulting plasmids were transformed into competent C41 *E. coli* cells for expression. A single colony from each transformation was picked and grown overnight at 37°C in 100 ml of 2xYT (16.0 g/L tryptone, 10.0 g/L yeast extract, 5.0 g/L NaCl) media containing 100 μg/ml of ampicillin. These cultures were then used to seed 1 L of 2xYT media. Cultures were induced at an OD600 of 0.9 with IPTG (0.5 mM final concentration), and grown for a further 4 hours at 37°C. The cells were harvested by centrifugation. Adnectin-anti-VEGFR2 and FN3Con-anti-VEGFR2 had their cell pellets resuspended in 5 ml/g of native lysis buffer (50 mM NaH_2_PO_4_, 300 mM NaCl, 10 mM imidazole, pH 8.0) and were lysed by sonication. Cell debris was removed by centrifugation and incubated in lysis buffer + 40 mM beta-mercaptoethanol to reduce disulfide bonds. Recombinant protein was then isolated from the supernatant by nickel affinity chromatography using loose Ni-NTA resin (Sigma). Protein eluted from Ni-NTA resin was filtered and then loaded onto a size exclusion column (Superdex 75 16/60, GE Healthcare) equilibrated in either PBS (140 mM NaCl, 2.7 mM KCl, 10 mM PO_4_^3-^, 4mM beta-mercaptoethanol pH 7.4) for biophysical characterization. Protein concentration was determined by Nanodrop ND-1000 (ThermoFisher) and protein was stored at 4 °C until use.

### Biotin-conjugation of proteins

Biotin was conjugated to lysines, which are on the non-binding loops of FN3Con-anti-VEGFR2 and Adnectin-anti-VEGFR2, to improve sensitivity and loading in ELISA or BLItz binding assays (EZ-Link™ Sulfo-NHS-LC-Biotinylation Kit, ThermoFisher 21435).

### Binding studies

#### SPR

The binding affinity of FN3Con-anti-VEGFR2 was measured using surface plasmon resonance with a 30 µl/min flow rate at 25 °C (BIAcore T-100, GE Healthcare). VEGFR2 domains (SiboBiological, 10012-H08H) were conjugated on a CM5 sensor chip through NHS/EDC activation and ethanolamine deactivation. HBS-EP (10 mM HEPES, 150 mM NaCl, 0.005% (v/v) Tween 20, 0.1% BSA pH 7.4) was used as the running buffer and FN3Con-anti-VEGFR2 was prepared in serial dilutions of HBS-EP. Multi-cycle kinetics were performed with 240 s association of FN3Con-anti-VEGFR2 and 800 s dissociation with blank HBS-EP buffer, followed by regeneration of the surface (30 s Glycine-HCl pH 1.5 followed by a 150 s stabilization period, then 30s NaOH pH 10 followed by a 300s stabilization period).

#### ELISA

96-well plates were coated overnight with 10 µg/ml VEGFR2, shaking at 4 °C. Wells were washed with PBS and blocked with milk blocking solution (5% powdered milk in PBS-T) for 2 hours. Wells were washed again and incubated with biotinylated anti-VEGFR2 proteins for 1 hour before washing and incubation with anti-biotin HRP solution for another hour. After a final wash, TMB was added to wells and left to incubate for up to 30 minutes. Fluorescence at 415-450 nm was read by a plate reader for each well.

### Binding calculations

High non-specific binding was a limiting factor in the analysis of affinity especially when attempting to determine kinetic constants which were compromised also by a large component of mass transport during association and ligand rebinding during dissociation. Consequently, the steady-state value for association of each FN3Con-anti-VEGFR2 concentration was determined at 190 s after injection, and a *K*_D_ was calculated using a plot of RU values at this point as a function of concentration with GraphPad Prism and a simple Langmuir type fit (Y=Bmax*X/(Kd+X) + NS*X + Background, where Y is the RU, X is the concentration in nM, and Background is constrained to 0).

The ELISA data was also limited by non-specific binding, however the aim of this experiment was to provide a comparison between Adnectin-anti-VEGFR2 and FN3Con-anti-VEGFR2. ELISA data was plotted in GraphPad Prism and a one site binding (hyperbola) model (Y= (B_max_ *X) / (K_d_ + X)) was applied to calculate the *K*_D_ as the midpoint of signal increase, this was presented as *K*_D_ alongside the R^2^ measure of fit. The Biacore analysis was presented as the most representative estimation of affinity, as the steady state approach allowed for better understanding of the effect of non-specific binding.

### Thermal stability

Thermal stability of purified Adnectin-anti-VEGFR2 and FN3Con-anti-VEGFR2 was measured by circular dichroism (CD). CD measurements were performed using a Jasco 815 spectropolarimeter with 0.2 mg/ml protein in PBS used in a 0.1 cm path length quartz cuvette. Thermal denaturation was measured by observing signal changes at 222 nm during heating at a rate of 1 °C/min. The melting temperature (*T*_m_) was obtained by fitting to a sigmoidal dose-response (variable slope) equation (Y=Bottom+(Top-Bottom)/(1+exp((V50-X)/Slope))).

### Long term stability (LTS)

Long term stability was measured by diluting purified protein to 1mg/ml in PBS + 2mM beta-mercaptoethanol pH 7.4, then storing in 1.7 ml Eppendorf tube in a temperature-controlled rooms at 4 °C, 21 °C and 36 °C. Stability was investigated as the size of monomer peak on a size exclusion column (Superdex 75 16/60, GE Healthcare) after injection of 100 µl volumes, 0.2 µm filtered, from each temperature fraction at 3, 6, 12, 22, and 24 months of storage. Peak areas were integrated by GE Unicorn software and percentages calculated from individual peak area divided by total area under the curve.

### Accelerated Stability

Accelerated stability was measured by diluting purified protein to 1mg/ml in 2 mM Beta-mercaptoethanol pH 7.4 with 62.4 mM of either Glycine, Histidine, Aspartic Acid, Arginine amino acids or Tween80, then storing in 1.7 ml Eppendorf tube in a heat block at 40 °C for 30 days. Stability was investigated as the size of monomer peak on a size exclusion column (Superdex 75 16/60, GE Healthcare) from injection of 500 µl volumes, 0.2 µm filtered, from each sample.

## Supplementary figures

**Figure S1:**
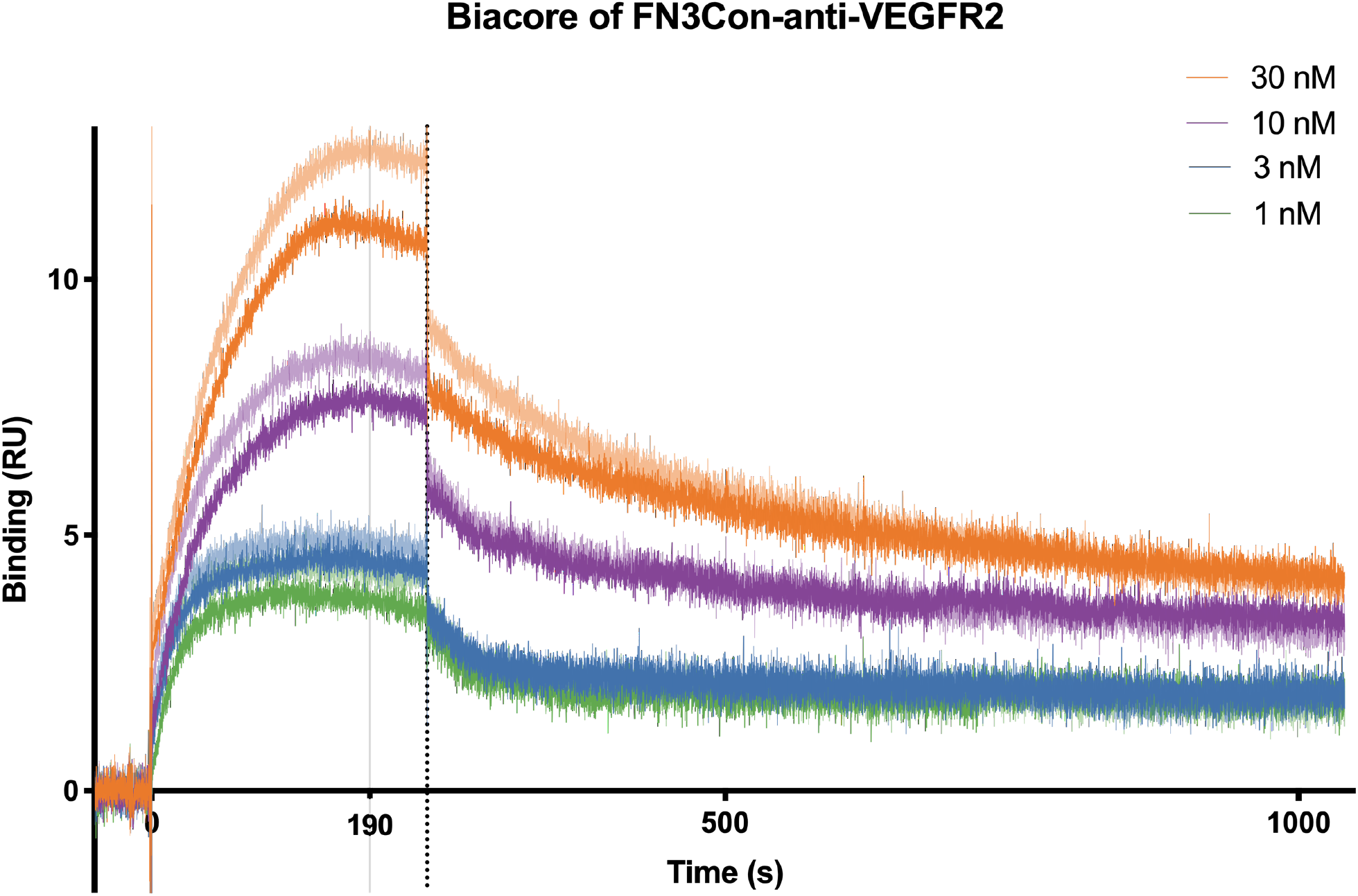
FN3Con-anti-VEGFR2 Biacore data

**Table S1:**
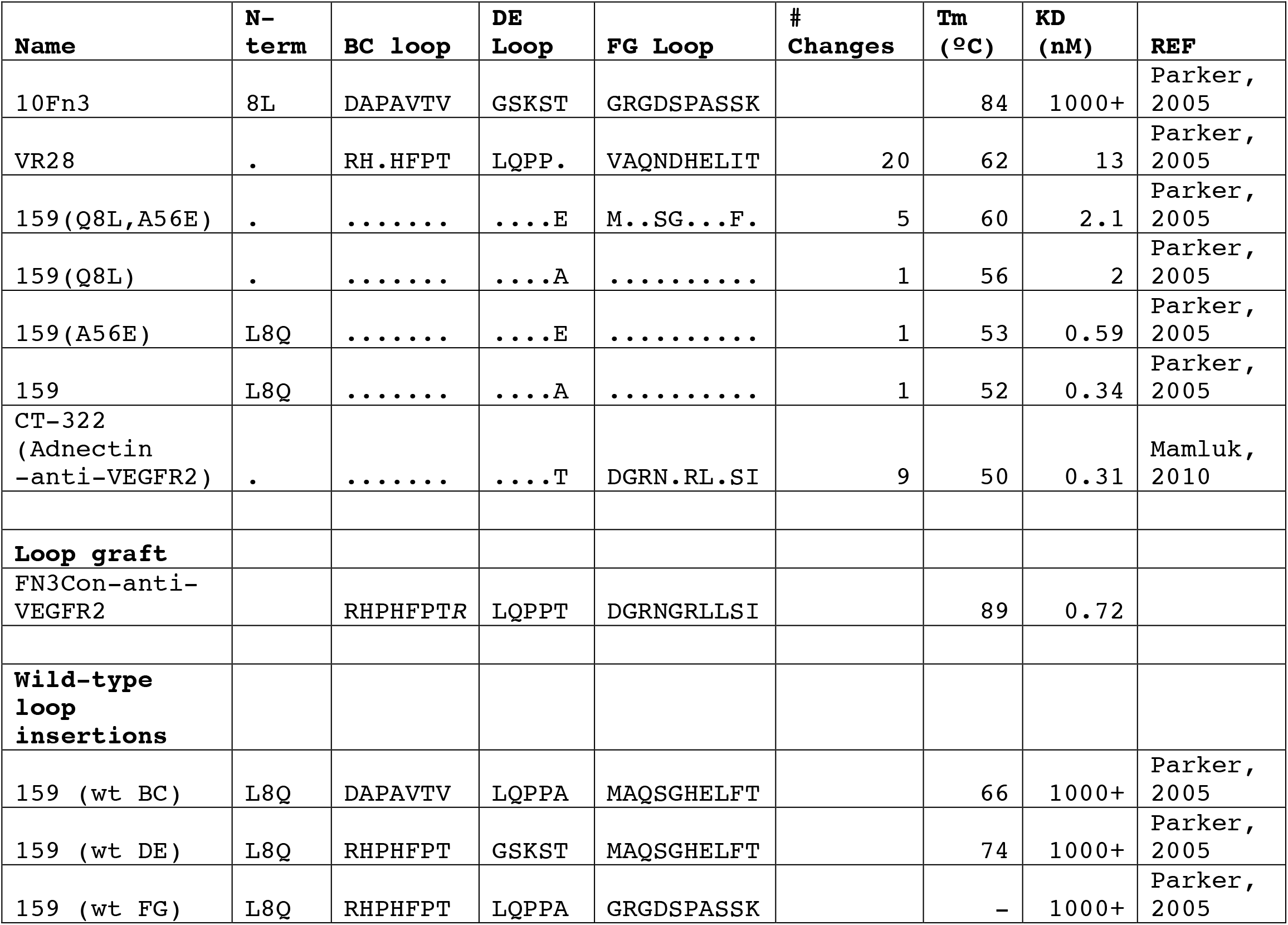
Table of loop sequence changes over the course of variation for function

## Notes

### Competing Interest Statement

The authors have declared no competing interest.

## References

1. Crook, Z. R., Nairn, N. W., and Olson, J. M. (2020) Miniproteins as a Powerful Modality in Drug Development. Trends in Biochemical Sciences. 45, 332–346

2. Vazquez-Lombardi, R., Phan, T. G., Zimmermann, C., Lowe, D., Jermutus, L., and Christ, D. (2015) Challenges and opportunities for non-antibody scaffold drugs. Drug Discovery Today. 20, 1271–1283

3. Koide, S., Koide, A., and Lipovsek, D. (2012) Target-binding proteins based on the 10th human fibronectin type III domain (10Fn3). Methods in Enzymology. 503, 135–156

4. Hantschel, O. (2017) Monobodies as possible next-generation protein therapeutics – a perspective. Swiss Medical Weekly. 10.4414/smw.2017.14545

5. Chandler, P. G., and Buckle, A. M. (2020) Development and Differentiation in Monobodies Based on the Fibronectin Type 3 Domain. Cells. 9, 610

6. Lipovsek, D. (2011) Adnectins: engineered target-binding protein therapeutics. Protein Engineering Design and Selection. 24, 3–9

7. Koide, A., Bailey, C. W., Huang, X., and Koide, S. (1998) The fibronectin type III domain as a scaffold for novel binding proteins. Journal of Molecular Biology. 284, 1141–1151

8. Stein, E., Toth, P., Butcher, M., Kereiakes, D., Magnu, P., Bays, H., Zhou, R., and Turner, T. A. (2019) Safety, Tolerability And Ldl-C Reduction With A Novel Anti-Pcsk9 Recombinant Fusion Protein (Lib003): Results Of A Randomized, Double-Blind, Placebo-Controlled, Phase 2 Study. Atherosclerosis. 287, e7

9. Wensel, D., Sun, Y., Davis, J., Li, Z., Zhang, S., McDonagh, T., Langley, D., Mitchell, T., Tabruyn, S., Nef, P., Cockett, M., and Krystal, M. (2019) GSK3732394: a Multi-specific Inhibitor of HIV Entry. Journal of Virology. 93, 1–21

10. Addis, R. C., Kolakowski, R., Kulkarni, S., Gorsky, J., Meyer, R., Xin, Y., Mortezavi, E., O’Neil, K. T., and Nadler, S. G. (2020) Abstract 1825: Tumor-targeted knockdown of KRAS mutants with novel Centyrin:siRNA conjugates. 10.1158/1538-7445.am2020-1825

11. Addis, R., Kolakowski, R., Kulkarni, S., Gorsky, J., Meyer, R., and ONeil, K. (2019) Abstract 4830: ABX9xx: A bispecific centyrin that synergizes to attenuate intracellular signaling in Met/EGFR positive tumors. in Experimental and Molecular Therapeutics, pp. 4830–4830, American Association for Cancer Research, 79, 4830–4830

12. Gregory, T. K., Berdeja, J. G., Patel, K. K., Ali, S. A., Cohen, A. D., Costello, C., Ostertag, E. M., Silva, N. de, Shedlock, D. J., Resler, M., Spear, M. A., and Orlowski, R. Z. (2018) Abstract CT130: Clinical trial of P-BCMA-101 T stem cell memory (Tscm) CAR-T cells in relapsed/refractory (r/r) multiple myeloma (MM). in Clinical Trials, pp. CT130–CT130, American Association for Cancer Research, 10.1158/1538-7445.am2018-ct130

13. Karnell, J. L., Albulescu, M., Drabic, S., Wang, L., Moate, R., Baca, M., Oganesyan, V., Gunsior, M., Thisted, T., Yan, L., Li, J., Xiong, X., Eck, S. C., De Los Reyes, M., Yusuf, I., Streicher, K., Müller-Ladner, U., Howe, D., Ettinger, R., Herbst, R., and Drappa, J. (2019) A CD40L-targeting protein reduces autoantibodies and improves disease activity in patients with autoimmunity. Science Translational Medicine. 10.1126/scitranslmed.aar6584

14. Löfblom, J., Frejd, F. Y., and Ståhl, S. (2011) Non-immunoglobulin based protein scaffolds. Current Opinion in Biotechnology. 22, 843–848

15. Olson, C. A., Liao, H. I., Sun, R., and Roberts, R. W. (2008) mRNA display selection of a high-affinity, modification-specific phospho-iκbα-binding fibronectin. ACS Chemical Biology. 3, 480–485

16. Jain, T., Sun, T., Durand, S., Hall, A., Houston, N. R., Nett, J. H., Sharkey, B., Bobrowicz, B., Caffry, I., Yu, Y., Cao, Y., Lynaugh, H., Brown, M., Baruah, H., Gray, L. T., Krauland, E. M., Xu, Y., Vásquez, M., and Wittrup, K. D. (2017) Biophysical properties of the clinical-stage antibody landscape. Proceedings of the National Academy of Sciences. 114, 944–949

17. Jarasch, A., Koll, H., Regula, J. T., Bader, M., Papadimitriou, A., and Kettenberger, H. (2015) Developability assessment during the selection of novel therapeutic antibodies. Journal of Pharmaceutical Sciences. 104, 1885–1898

18. Chandler, P. G., Broendum, S. S., Riley, B. T., Spence, M. A., Jackson, C. J., McGowan, S., and Buckle, A. M. (2020) Strategies for Increasing Protein Stability. in Methods in Molecular Biology, pp. 163–181, 10.1007/978-1-4939-9869-2_10

19. Tokuriki, N., Stricher, F., Schymkowitz, J., Serrano, L., and Tawfik, D. S. (2007) The Stability Effects of Protein Mutations Appear to be Universally Distributed. Journal of Molecular Biology. 369, 1318–1332

20. Tokuriki, N., and Tawfik, D. S. (2009) Stability effects of mutations and protein evolvability. Current Opinion in Structural Biology. 19, 596–604

21. Bloom, J. D., Labthavikul, S. T., Otey, C. R., and Arnold, F. H. (2006) Protein stability promotes evolvability. Proceedings of the National Academy of Sciences. 103, 5869–5874

22. Besenmatter, W., Kast, P., and Hilvert, D. (2006) Relative tolerance of mesostable and thermostable protein homologs to extensive mutation. Proteins: Structure, Function, and Bioinformatics. 66, 500–506

23. Romero, P. A., and Arnold, F. H. (2009) Exploring protein fitness landscapes by directed evolution. Nature Reviews Molecular Cell Biology. 10, 866–876

24. Ota, N., Kurahashi, R., Sano, S., and Takano, K. (2018) The direction of protein evolution is destined by the stability. Biochimie. 150, 100–109

25. Falconer, R. J. (2019) Advances in liquid formulations of parenteral therapeutic proteins. Biotechnology Advances. 37, 107412

26. Falconer, R. J., Chan, C., Hughes, K., and Munro, T. P. (2011) Stabilization of a monoclonal antibody during purification and formulation by addition of basic amino acid excipients. Journal of Chemical Technology and Biotechnology. 86, 942–948

27. Berteau, C., Filipe-Santos, O., Wang, T., Roja, H. E., Granger, C., and Schwarzenbach, F. (2015) Evaluation of the impact of viscosity, injection volume, and injection flow rate on subcutaneous injection tolerance. Medical Devices: Evidence and Research. 8, 473–484

28. Wang, W. (2015) Tolerability of hypertonic injectables. International Journal of Pharmaceutics. 490, 308–315

29. Zbacnik, T. J., Holcomb, R. E., Katayama, D. S., Murphy, B. M., Payne, R. W., Coccaro, R. C., Evans, G. J., Matsuura, J. E., Henry, C. S., and Manning, M. C. (2017) Role of Buffers in Protein Formulations. Journal of Pharmaceutical Sciences. 106, 713–733

30. Wilding, M., Hong, N., Spence, M., Buckle, A. M., and Jackson, C. J. (2019) Protein engineering: The potential of remote mutations. Biochemical Society Transactions. 47, 701–711

31. Porebski, B. T., and Buckle, A. M. (2016) Consensus protein design. Protein Engineering, Design and Selection. 29, 245–251

32. Kumar, S., and Nussinov, R. (2001) How do thermophilic proteins deal with heat? Cellular and Molecular Life Sciences. 58, 1216–1233

33. Goldenzweig, A., and Fleishman, S. (2018) Principles of Protein Stability and Their Application in Computational Design. 10.1146/annurev-biochem

34. Sternke, M., Tripp, K. W., and Barrick, D. (2019) Consensus sequence design as a general strategy to create hyperstable, biologically active proteins. Proceedings of the National Academy of Sciences of the United States of America. 166, 11275–11284

35. Porebski, B. T., Nickson, A. A., Hoke, D. E., Hunter, M. R., Zhu, L., McGowan, S., Webb, G. I., and Buckle, A. M. (2015) Structural and dynamic properties that govern the stability of an engineered fibronectin type III domain. Protein Engineering, Design and Selection. 28, 67–78

36. Porebski, B. T., Conroy, P. J., Drinkwater, N., Schofield, P., Vazquez-Lombardi, R., Hunter, M. R., Hoke, D. E., Christ, D., McGowan, S., and Buckle, A. M. (2016) Circumventing the stability-function trade-off in an engineered FN3 domain. Protein Engineering Design and Selection. 29, 1–9

37. Parker, M. H., Chen, Y., Danehy, F., Dufu, K., Ekstrom, J., Getmanova, E. V., Gokemeijer, J., Xu, L., and Lipovsek, D. (2005) Antibody mimics based on human fibronectin type three domain engineered for thermostability and high-affinity binding to vascular endothelial growth factor receptor two. Protein Engineering, Design and Selection. 18, 435–444

38. Mamluk, R., Carvajal, I. M., Morse, B. A., Wong, H. K., Abramowitz, J., Aslanian, S., Lim, A.-C., Gokemeijer, J., Storek, M. J., Lee, J., Gosselin, M., Wright, M. C., Camphausen, R. T., Wang, J., Chen, Y., Miller, K., Sanders, K., Short, S., Sperinde, J., Prasad, G., Williams, S., Kerbel, R. S., Ebos, J., Mutsaers, A., Mendlein, J. D., Harris, A. S., and Furfine, E. S. (2010) Anti-tumor effect of CT-322 as an Adnectin inhibitor of vascular endothelial growth factor receptor-2. mAbs. 2, 199–208

39. Agarkhed, M., O’Dell, C., Hsieh, M. C., Zhang, J., Goldstein, J., and Srivastava, A. (2013) Effect of Polysorbate 80 concentration on thermal and photostability of a monoclonal antibody. AAPS PharmSciTech. 14, 1–9

40. Dineen, S. P., Sullivan, L. a, Beck, A. W., Miller, A. F., Carbon, J. G., Mamluk, R., Wong, H., and Brekken, R. a (2008) The Adnectin CT-322 is a novel VEGF receptor 2 inhibitor that decreases tumor burden in an orthotopic mouse model of pancreatic cancer. BMC cancer. 8, 352

41. Ackermann, M., Carvajal, I. M., Morse, B. A., Moreta, M., O’Neil, S., Kossodo, S., Peterson, J. D., Delventhal, V., Marsh, H. N., Furfine, E. S., and Konerding, M. A. (2010) Adnectin CT-322 inhibits tumor growth and affects microvascular architecture and function in Colo205 tumor xenografts. International Journal of Oncology. 38, 267–277

42. Waterman, P., Palanichamy, K., Morse, B., Gonda, D. D., Akers, J., Sanchez, C., Carvajal, I., Marsh, N., Waters, J. D., Furfine, E., Chen, C. C., Scheer, J. K., Sahin, A., Weissleder, R., Futalan, D., Carter, B. S., and Chakravarti, A. (2012) CT322, a VEGFR-2 antagonist, demonstrates anti-glioma efficacy in orthotopic brain tumor model as a single agent or in combination with temozolomide and radiation therapy. Journal of Neuro-Oncology. 110, 37–48

43. Ackermann, M., Morse, B. A., Delventhal, V., Carvajal, I. M., and Konerding, M. A. (2012) Anti-VEGFR2 and anti-IGF-1R-Adnectins inhibit Ewing’s sarcoma A673-xenograft growth and normalize tumor vascular architecture. Angiogenesis. 15, 685–695

44. Tolcher, A. W., Sweeney, C. J., Papadopoulos, K., Patnaik, A., Chiorean, E. G., Mita, A. C., Sankhala, K., Furfine, E., Gokemeijer, J., Iacono, L., Eaton, C., Silver, B. A., and Mita, M. (2011) Phase I and pharmacokinetic study of CT-322 (BMS-844203), a targeted adnectin inhibitor of VEGFR-2 based on a domain of human fibronectin. Clinical Cancer Research. 17, 363–371

45. Sachdev, E., Gong, J., Rimel, B., and Mita, M. (2015) Adnectin-Targeted Inhibitors: Rationale and Results. Current Oncology Reports. 17, 1–6

46. Schiff, D., Kesari, S., De Groot, J., Mikkelsen, T., Drappatz, J., Coyle, T., Fichtel, L., Silver, B., Walters, I., and Reardon, D. (2015) Phase 2 study of CT-322, a targeted biologic inhibitor of VEGFR-2 based on a domain of human fibronectin, in recurrent glioblastoma. Investigational New Drugs. 33, 247–253

47. Getmanova, E. V., Chen, Y., Bloom, L., Gokemeijer, J., Shamah, S., Warikoo, V., Wang, J., Ling, V., and Sun, L. (2006) Antagonists to Human and Mouse Vascular Endothelial Growth Factor Receptor 2 Generated by Directed Protein Evolution In Vitro. Chemistry and Biology. 13, 549–556

48. Aghaabdollahian, S., Cohan, R. A., Norouzian, D., and Davami, F. (2019) Enhancing bioactivity, physicochemical, and pharmacokinetic properties of a through PASylation technology. 10.1038/s41598-019-39776-0

49. Verhoeyen, M., Milstein, C., and Winter, G. (1988) Reshaping human antibodies: grafting an antilysozyme activity. Science. 239, 1534–1536

50. Ewert, S., Honegger, A., and Plückthun, A. (2004) Stability improvement of antibodies for extracellular and intracellular applications: CDR grafting to stable frameworks and structure-based framework engineering. Methods (San Diego, Calif.). 34, 184–99

51. Harrison, R. E., Man, C.-W., and Wang, Y. (2020) Abstract 598: Integrated computational and experimental design of a monobody targeting PDL1. 80, 598–598

52. Kadonosono, T., Yimchuen, W., Ota, Y., See, K., Furuta, T., Shiozawa, T., Kitazawa, M., Goto, Y., Patil, A., Kuchimaru, T., and Kizaka-Kondoh, S. (2020) Design Strategy to Create Antibody Mimetics Harbouring Immobilised Complementarity Determining Region Peptides for Practical Use. Scientific Reports. 10, 1–11

53. See, K., Kadonosono, T., Ota, Y., Miyamoto, K., Yimchuen, W., and Kizaka-Kondoh, S. (2020) Reconstitution of an Anti-HER2 Antibody Paratope by Grafting Dual CDR-Derived Peptides onto a Small Protein Scaffold. Biotechnology Journal. 15, 2000078

54. Natarajan, A., Patel, C. B., Ramakrishnan, S., Panesar, P. S., Long, S. R., and Gambhir, S. S. (2019) A Novel Engineered Small Protein for Positron Emission Tomography Imaging of Human Programmed Death Ligand-1: Validation in Mouse Models and Human Cancer Tissues. Clinical Cancer Research. 25, 1774–1785

55. Deonarain, M. P., and Yahioglu, G. (2021) Current strategies for the discovery and bioconjugation of smaller, targetable drug conjugates tailored for solid tumour therapy. Expert Opinion on Drug Discovery. 10.1080/17460441.2021.1858050

56. Plückthun, A. (2015) Designed Ankyrin Repeat Proteins (DARPins): Binding Proteins for Research, Diagnostics, and Therapy. Annual Review of Pharmacology and Toxicology. 55, 489– 511

57. Chen, T. F., de Picciotto, S., Hackel, B. J., and Wittrup, K. D. (2013) Engineering fibronectin-based binding proteins by yeast surface display. Methods in enzymology. 523, 303–26

58. Pike, R. M., and Chandler, C. H. (1972) Effect of storage on the ability of 2-mercaptoethanol to inactivate M antibody. Infection and immunity. 5, 416–417

59. Tokuriki, N., and Tawfik, D. S. (2009) Protein Dynamism and Evolvability. Science. 324, 203–207

60. Katayama, D. S., Nayar, R., Chou, D. K., Valente, J. J., Cooper, J., Henry, C. S., Vander Velde, D. G., Villarete, L., Liu, C. P., and Manning, M. C. (2006) Effect of buffer species on the thermally induced aggregation of interferon-tau. Journal of Pharmaceutical Sciences. 95, 1212– 1226

61. Jorgensen, L., Hostrup, S., Moeller, E. H., and Grohganz, H. (2009) Recent trends in stabilising peptides and proteins in pharmaceutical formulation - Considerations in the choice of excipients. Expert Opinion on Drug Delivery. 6, 1219–1230

62. Ota, C., Fukuda, Y., Tanaka, S., and Takano, K. (2020) Spectroscopic Evidence of the Salt-Induced Conformational Change around the Localized Electric Charges on the Protein Surface of Fibronectin Type III. Langmuir. 10.1021/acs.langmuir.0c02367

63. Platts, L., and Falconer, R. J. (2015) Controlling protein stability: Mechanisms revealed using formulations of arginine, glycine and guanidinium HCl with three globular proteins. International Journal of Pharmaceutics. 486, 131–135

64. Arakawa, T., and Tsumoto, K. (2003) The effects of arginine on refolding of aggregated proteins: Not facilitate refolding, but suppress aggregation. Biochemical and Biophysical Research Communications. 304, 148–152

65. Shukla, D., and Trout, B. L. (2010) Interaction of arginine with proteins and the mechanism by which it inhibits aggregation. Journal of Physical Chemistry B. 114, 13426–13438

66. Porebski, B. T., Keleher, S., Hollins, J. J., Nickson, A. A., Marijanovic, E. M., Borg, N. A., Costa, M. G. S., Pearce, M. A., Dai, W., Zhu, L., Irving, J. A., Hoke, D. E., Kass, I., Whisstock, J. C., Bottomley, S. P., Webb, G. I., McGowan, S., and Buckle, A. M. (2016) Smoothing a rugged protein folding landscape by sequence-based redesign. Scientific Reports. 6, 1–14

67. Chen, B., Bautista, R., Yu, K., Zapata, G. A., Mulkerrin, M. G., and Chamow, S. M. (2003) Influence of Histidine on the Stability and Physical Properties of a Fully Human Antibody in Aqueous and Solid Forms. Pharmaceutical Research. 20, 1952–1960

68. Asghar, W., Yuksekkaya, M., Shafiee, H., Zhang, M., Ozen, M. O., Inci, F., Kocakulak, M., and Demirci, U. (2016) Engineering long shelf life multi-layer biologically active surfaces on microfluidic devices for point of care applications. Scientific Reports. 6, 1–10

